# Beyond orphaned infants: novel effects of maternal death in wild primates

**DOI:** 10.1101/2020.07.22.212605

**Authors:** Matthew N. Zipple, Jeanne Altmann, Fernando A. Campos, Marina Cords, Linda M. Fedigan, Richard R. Lawler, Elizabeth V. Lonsdorf, Susan Perry, Anne E. Pusey, Tara S. Stoinski, Karen B. Strier, Susan C. Alberts

**Affiliations:** Duke University; Princeton University; University of Texas at San Antonio; Columbia University; University of Calgary; Boston University; Franklin & Marshall College; University of California, Los Angeles; Dian Fossey Gorilla Fund; University of Wisconsin-Madison

## Abstract

Primate offspring often depend on their mothers well beyond the age of weaning, and offspring that experience maternal death in early life can suffer substantial reductions in fitness across the lifespan. Here we leverage data from eight wild primate populations (seven species) to examine two underappreciated pathways linking early maternal death and offspring fitness that are distinct from direct effects of orphaning on offspring survival. First, we show that, for five of the seven species, offspring face reduced survival during the years immediately *preceding* maternal death, while the mother is still alive. Second, we identify an intergenerational effect of early maternal loss in three species (muriquis, baboons, and blue monkeys), such that early maternal death experienced in one generation leads to reduced offspring survival in the next. Our results have important implications for the evolution of slow life histories in primates, as they suggest that maternal condition and survival are more important for offspring fitness than previously realized.

## Introduction

Mammalian life history is marked by a strong dependent relationship between offspring and their mothers (1). The quantity or quality of maternal allocation to offspring, particularly during the gestation and lactation periods, is often related to maternal physical condition, and a range of offspring fitness outcomes are compromised if gestating or lactating mothers are in poor condition (2–8). In addition, infants that experience maternal loss prior to weaning face an enormous, acute risk of death in both non-human mammals and in humans (9–13) (Figure 1, blue arrow).

**Figure 1.**
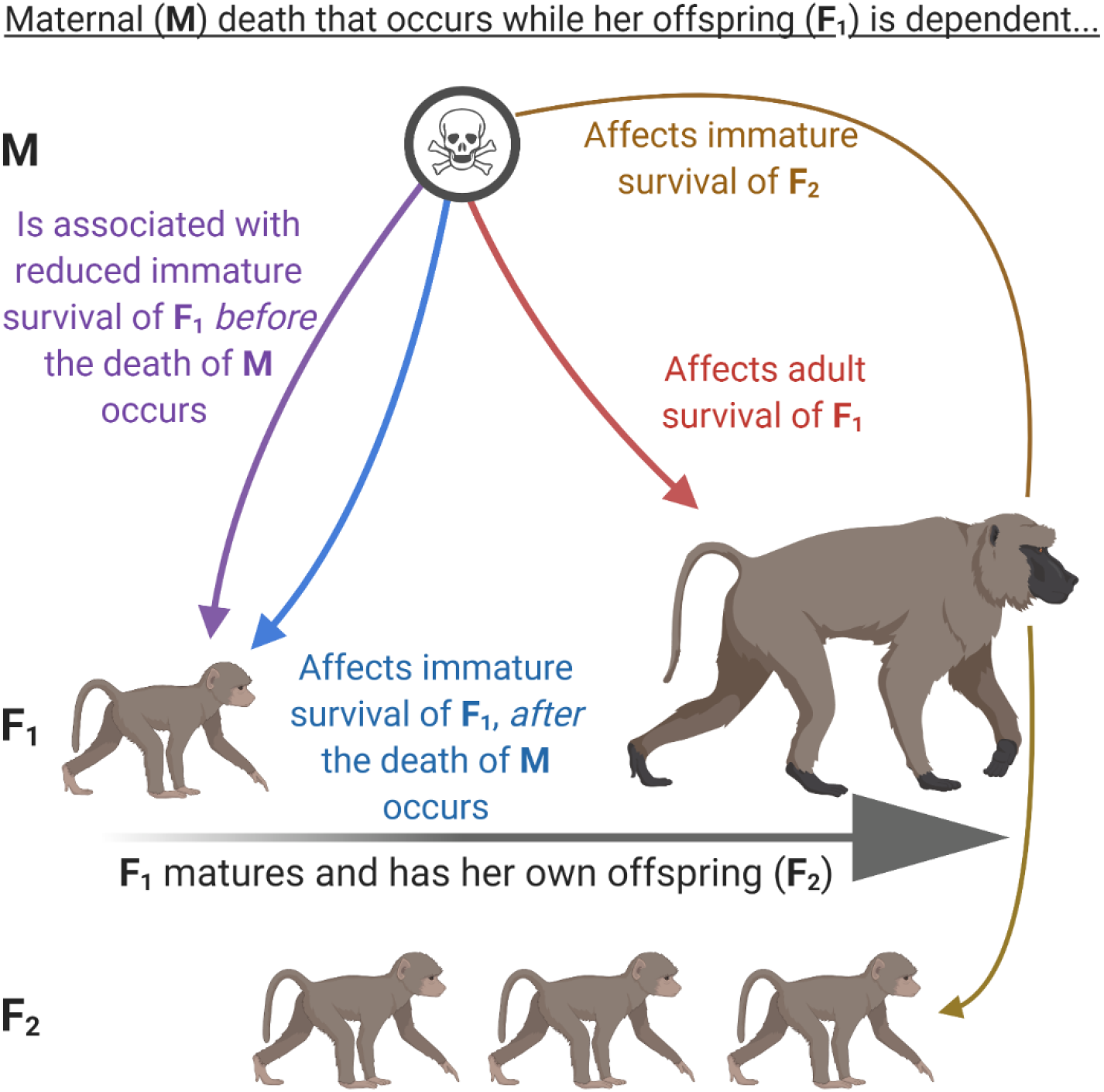
Four ways in which the death of a female primate mother (M) may be linked to her offspring’s fitness (F_1_), if the death of M occurs while F_1_ is still dependent on M. First, F_1_ should display reduced survival during the immature period, *following* the death of M (especially before weaning but also after), because F_1_ will lack the critically important social, nutritional, and/or protective resources that M provided (blue arrow). Second, F_1_ should display reduced survival in the period *before* M actually dies, because, on average, mothers are in worse condition shortly before their death compared to mothers that survive the same period. We therefore expect M to provide lower quality maternal care to F_1_ during the weeks to years immediately preceding M’s death (purple arrow). Third, if F_1_ survives these first two challenges, she is likely to be in chronically worse condition during adulthood because of reductions in maternal allocation that she received during development (red arrow). F_1_ should therefore face reduced survival in adulthood, years (or even decades) after the death of M occurred. Fourth, this chronic reduction in F_1_’s condition may have an intergenerational effect, such that F_2_ (F_1_’s offspring) also experience reduced immature survival (gold arrow). The analyses presented here focus on the purple arrow and the gold arrow.

In some species, including primates, hyenas, whales, and some ungulates, mothers and offspring continue to associate after weaning, and mothers may provide substantial social and energetic input as well as protection during some or all of the remainder of the pre-dispersal, immature period (hereafter the ‘immature period’) (12, 14–21). Thus, loss of the mother can continue to heighten the risk of death even in weaned, immature offspring (17, 19, 21) (Figure 1, blue arrow). However, because offspring are less dependent on mothers during this phase of life, the effects of maternal loss after weaning can be sub-lethal (9, 16, 22). If an offspring that is weaned (but still partially dependent on its mother) survives its mother’s death, the offspring may experience long-lasting negative effects, including adverse behavioral or social outcomes in adolescence or adulthood (humans: (23, 24); non-humans: (19, 20, 25–31)). In baboons, chimpanzees, and elephants, motherless offspring may experience reduced survival during adolescence and adulthood, well after the maternal loss occurs, presumably because maternal loss results in a chronic reduction in body condition (13, 21, 26, 32) (Figure 1, red arrow).

These observations (see citations in previous paragraphs) combine to provide a strong framework that describes the dependent relationship between mammalian offspring and their mothers and allows us to make predictions about the expected effects of maternal death on offspring fitness. This framework includes the following observations and conjectures: Mammalian offspring are critically dependent on their mothers for nutrition, protection, transport, and learning. In many species, the period of dependence is not restricted to infancy and may extend well past weaning. Poor maternal body condition during this dependent period can lead to reduced maternal allocation to offspring and hence poor offspring body condition, which may have both immediate and later-life consequences for offspring fitness outcomes, including survival. As a result, mothers in poor condition (even those that survive to wean their offspring) are likely to produce offspring in poor condition that experience compromised survival. Maternal death at any time during this dependent period can therefore result in both short-term and chronic reductions in offspring physical condition and survival.

This framework yields four main predictions about how maternal death experienced in early life affects offspring fitness outcomes across the lifespan. First, immature offspring that lose their mother will face reduced survival throughout the remainder of their immature period (Figure 1, blue arrow). Although the impact of maternal loss is expected to be especially strong if the mother dies before the offspring is weaned, the immature offspring may continue to face reduced survival if its mother dies any time before the offspring matures. Second, because the loss of the mother in early life results in developmental constraints that persist throughout the offspring’s lifetime, offspring that experienced early maternal loss will continue to experience reduced survival in adulthood, leading to shortened adult lifespans (Figure 1, red arrow). These two predictions are important for offspring fitness outcomes; they have been previously tested in several species (see above summary) and are therefore not the focus of our study.

We focus instead on two additional predictions, which have received little previous attention. First, we expect offspring to face reduced survival if their mothers are going to die in the near future, because, on average, a mother whose death is imminent is more likely to be in poor condition as compared to those mothers that survive the same period. We can test this prediction by measuring the association between offspring survival and impending maternal death, *while the mother is still alive* (Figure 1, purple arrow). Second, we predict an intergenerational effect of early maternal loss on offspring survival (Figure 1, gold arrow). That is, we predict that female offspring that experience maternal loss but still survive to adulthood (F_1_ generation in Figure 1) will produce offspring with compromised survival (F_2_ generation). We expect the proximate mechanism leading to this intergenerational effect to be that F_1_’s compromised condition causes her to be less able to allocate adequate resources to her offspring.

These latter two predictions about survival patterns have been previously confirmed in wild baboons(33), but otherwise we have little knowledge of the generality of these two links between maternal survival and the survival of offspring (F_1_ generation) and grand offspring (F_2_ generation) in natural populations of primates or other mammals. Here we leverage long-term longitudinal data from eight wild populations of seven primate species to assess (1) the extent to which offspring suffer reduced survival when their mothers will soon die (Figure 1, purple arrow), and (2) the extent to which the effects of early maternal loss carry over from one generation to the next, resulting in reduced immature survival for offspring whose mothers experienced early maternal loss (Figure 1, gold arrow).

## Results

The Primate Life Histories Database (PLHD) is a collection of demographic data from seven primate species across eight long-term studies of wild populations. The PLHD contains data from taxonomically diverse primates, including great apes (eastern chimpanzees [*Pan troglodytes schweinfurthii*] and mountain gorillas [*Gorilla beringei beringei*]), Old World monkeys (yellow baboons [*Papio cynocephalus*] and blue monkeys [*Cercopithecus mitis*]), New World monkeys (northern muriquis [*Brachyteles hypoxanthus*] and two populations of white-faced capuchins [*Cebus capucinus imitator*]), and indriids (Verreaux’s sifakas [*Propithecus verreauxi*]). Each of the studies and the database have been described elsewhere (34–36). Of critical importance for the present study, data are available on the timing of major life history events, including birth and death dates for individuals from all species. Sample sizes ranged from 71 to 1123 offspring from a given population, depending on the analysis and on the eligibility of offspring for inclusion (see Methods, Table S1).

### *First prediction*: offspring are more likely to die if their mothers face impending death

To measure the association between offspring survival and impending maternal death, we built mixed effects Cox proportional hazards models of offspring survival during the first two years of life. The predictor of interest is a binary fixed effect indicator of whether the mother died within four years after the offspring’s birth. This predictor represents our estimate of impending maternal death. We also included random effects of maternal ID and site-specific birth year to control for cohort effects in particular years in each study site (see Methods). For offspring whose mothers died before the offspring or on the same day as the offspring, we right-censored the offspring’s survival at the day of maternal death. That is, we included in the model of offspring survival only those periods of the offspring’s first two years of life when mothers were alive; the model did not consider offspring survival after maternal death occurred, because we were only interested in determining whether offspring faced reduced survival in the period *before* impending maternal death.

We built species-specific models using data from each of the seven species. Each species is represented in the database by a single population, except for capuchins, which are studied both at Santa Rosa (SR) and Lomas Barbudal (LB). When modelling capuchin survival, we therefore built both population-specific models and a combined-population model that additionally included a fixed effect of study site ID. Finally, we built a combined species model that considered data from all seven species together and a model that combined data from all species except baboons, which made up about 1/3 of all records. In addition to maternal ID and site-specific birth year, these combined-species models also included a random effect of study site ID.

In our species-level models, five species showed statistically significant associations between offspring survival during years 0-2 and impending maternal death. Specifically, offspring of muriquis (Hazard Ratio [HR] = 2.24, z = 2.24, p = 0.03), capuchins (HR=1.65, z = 2.64, p = 0.008), chimpanzees (HR = 2.18, z = 2.49, p = 0.01), baboons (HR = 1.34, z = 2.17, p = 0.03), and sifakas (HR = 1.33, z = 2.07, p = 0.045) were less likely to survive their first two years of life if their mothers were going to die in the near future (note that a hazard ratio greater than 1 indicates an *increase* in mortality and hence a *reduction* in survival). The other two species (gorillas and blue monkeys) did not show statistically significant associations when considered on their own, but the coefficient estimates were both in the expected direction (Table 1).

**Table 1.** Results of mixed effects Cox proportional hazards models that predict offspring survival in years 0-2 as a function of impending maternal death. All models include site-specific random effects of birth year and random effects of maternal ID. Bold values refer to a statistically significant effect (‘p value’ < 0.05) or an estimate in the expected direction (‘In expected direction?’ is ‘Yes’).

| Species | Coef. Estimate | Std. Error | z value | p value | Hazard Ratio Est. [95% Conf. Int.] | In expected direction? |
| --- | --- | --- | --- | --- | --- | --- |
| All Species Combined<sup>^</sup> | 0.35 | 0.08 | 4.54 | **<0.0001** | 1.42<br>[1.22,1.65] | Yes |
| All Species, Except Baboons<sup>^</sup> | 0.39 | 0.09 | 4.16 | **<0.0001** | 1.47<br>[1.22,1.76] | Yes |
| Muriquis | 0.81 | 0.36 | 2.24 | **0.03** | 2.24<br>[1.11,4.55] | Yes |
| Chimpanzees | 0.78 | 0.32 | 2.49 | **0.01** | 2.18<br>[1.17,4.08] | Yes |
| Capuchins (Santa Rosa) | 0.45 | 0.30 | 1.49 | 0.14 | 1.57<br>[0.87,2.82] | Yes |
| Capuchins (Lomas Barbudal) | 0.57 | 0.25 | 2.29 | **0.02** | 1.77<br>[1.08,2.89] | Yes |
| Capuchins (Combined)* | 0.50 | 0.19 | 2.64 | **0.008** | 1.65<br>[1.14,2.39] | Yes |
| Baboons | 0.29 | 0.13 | 2.17 | **0.03** | 1.34<br>[1.04,1.72] | Yes |
| Sifakas | 0.28 | 0.14 | 2.01 | **0.045** | 1.33<br>[1.01,1.76] | Yes |
| Gorillas | 0.29 | 0.53 | 0.55 | 0.59 | 1.34<br>[0.47,3.78] | Yes |
| Blue Monkeys | 0.08 | 0.22 | 0.38 | 0.70 | 1.08<br>[0.70,1.67] | Yes |

<sup>^</sup>Model also included random effect of study ID.
\*Model also included fixed effect of study ID.

When we built a single model with data from all species, the association between offspring survival and impending maternal death was strong and highly statistically significant (HR = 1.41, z = 4.59, p < 0.0001). This association remained strong and significant even when we excluded baboons (the largest sample size in the database) from the analysis (HR = 1.46, z = 4.21, p < 0.0001; Figure 2, Table 1, Figure S1).

**Figure 2.**
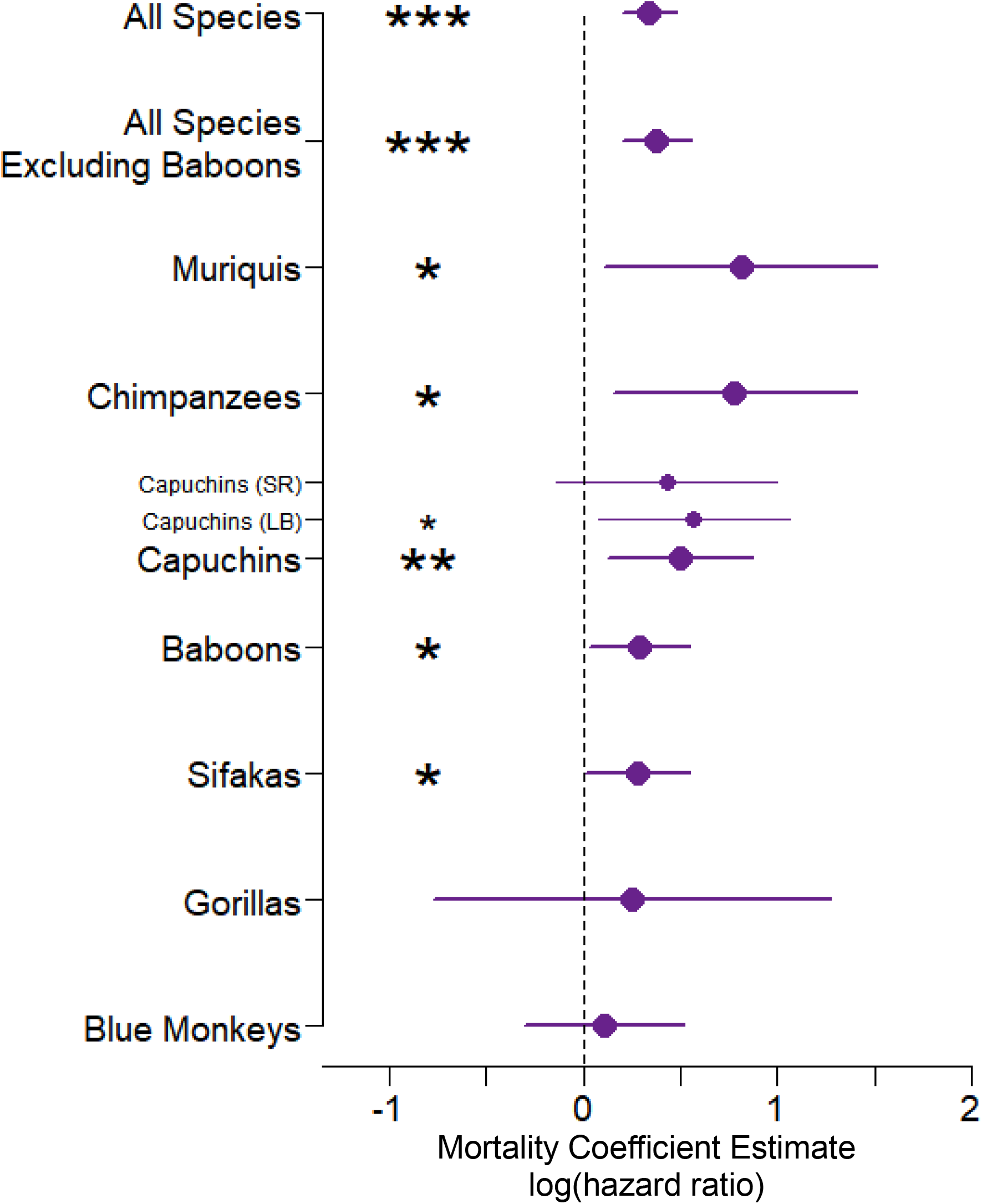
Impending maternal death predicts offspring survival in 5 of 7 primate species, and in combined-species models. Closed circles show, for each primate population, the magnitude and direction of the mortality coefficient estimates (equivalent to the log of the hazard ratio) for the association between offspring mortality (from birth to 2 years) and impending maternal death, with 95% confidence intervals. The vertical dashed line at a coefficient estimate of zero indicates no effect, with positive values indicating *increased* mortality and hence *reduced survival*. Significance levels are indicated as follows: (*** p < 0.001, ** p < 0.01, * p ≤ 0.05).

Our definition of offspring deaths linked to impending maternal death included offspring that died as neonates but whose mothers did not die until nearly 4 years later. Is it reasonable to expect an association to exist between infant death and maternal death when the events are separated by such long periods? To examine this question, we conducted an additional analysis considering offspring survival during the first six months of life only, and including only offspring whose mothers either (1) survived for the entire 4 years following offspring birth or (2) died 3.5 to 4 years after offspring birth, i.e., quite a long time after offspring birth. In a combined-species model of offspring survival, our single predictor was an indicator of whether the mother would die during the period 3.5 to 4 years after offspring birth). Remarkably, we found a strong and significant association between offspring survival in the first 6 months of life and maternal death in years 3.5 to 4 after offspring birth (HR [95% CI] = 1.7 [1.1,2.6]; p = 0.02). In other words, young infants are more likely to die if their mothers are going to die *more than three years later* (Figure S3). This result suggests that primate offspring may be quite sensitive to maternal condition.

A possible explanation for the negative association between immature survival and impending maternal death is that offspring may have died as a result of being born to old mothers, as older mothers may have been more likely to die in the near future. To test this possibility, we added mothers’ chronological ages to the combined-species model, as well as to the species-level models for baboons, chimpanzees, muriquis, capuchins, and sifakas. We used z-scored chronological age for each study site because of differences in life expectancy across studies. In no cases did we identify a significant effect of mothers’ chronological age on offspring survival after controlling for the timing of maternal death relative to offspring birth. The models for chimpanzee, capuchin, baboon and combined species all continued to indicate a statistically significant effect of impending maternal death on offspring survival after controlling for maternal chronological age (see Table S2), and although the coefficient estimates in muriquis and sifakas were no longer significant with the addition of maternal age, the estimated effect of impending maternal death on offspring survival was little changed by the addition of maternal chronological age (muriquis: HR = 2.05 with and HR = 2.24 without; sifakas: HR = 1.29 with and HR = 1.33 without).

Our results suggest that females are, on average, less able to keep their offspring alive as they get older and approach death. However, this effect occurs regardless of the *chronological* age of the mother: an offspring that is born to a mother who will soon die is at an elevated risk of death regardless of whether that mother is old or young. This result suggests that maternal senescence does play a critical role in offspring outcomes, but that senescence begins at different ages or proceeds at different rates in different mothers (see also (37), Figure 3, for related analyses on baboons).

**Figure 3.**
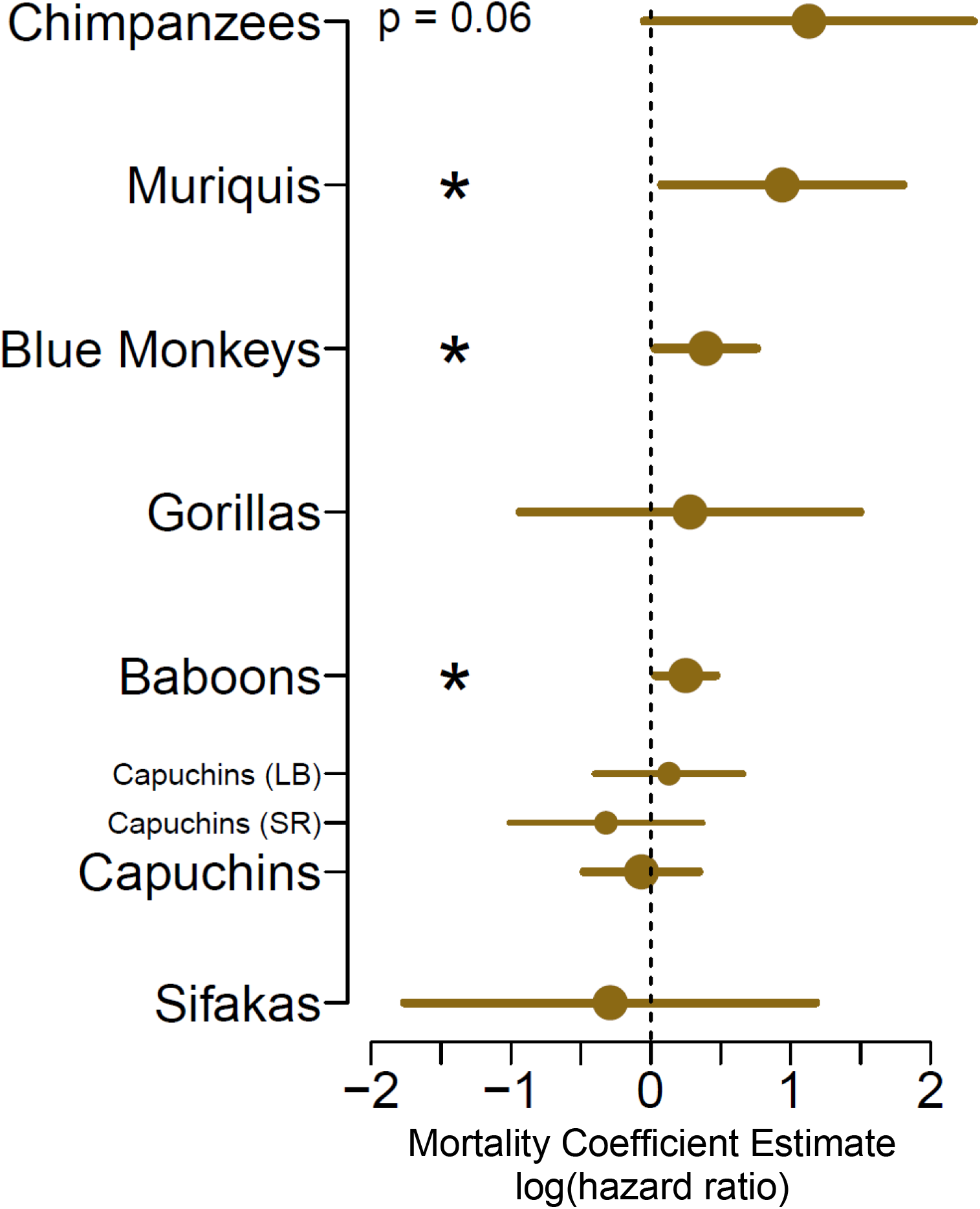
Maternal loss experienced by F1 mothers results in increased mortality for F2 offspring. Closed circles show, for each primate population, the magnitude and direction of the mortality coefficient estimates (equivalent to the log of the hazard ratio) for the association between F2 offspring mortality and maternal loss experienced by the F1 generation, 95% confidence intervals. The vertical dashed line at a coefficient estimate of zero indicates no effect, with positive values indicating *increased* mortality and hence *reduced survival*. An asterisk indicates a p value < 0.05.

Another possible explanation for the negative association between immature survival and impending maternal death is that this association is driven by covariation in the offspring’s and the mother’s extrinsic mortality risk. This could occur, for instance, if the social groups that they belong to experience higher predation pressure or occupy lower quality home ranges as compared to other social groups. To test this possibility, we performed a post hoc analysis in which we replaced the random effect of study site-specific birth year with a random effect of social group-specific birth year (each study site has within it multiple social groups). This change did not result in any substantial changes to the coefficient estimates of the effect of impending maternal death in any models, and did not affect qualitative results for 7 of the 8 populations. This change did cause the p value associated with the sifaka model to increase from 0.045 to 0.051, but the magnitude of the coefficient estimate for the sifaka model actually *increased* slightly (0.29 vs 0.28, see Table S3 for full results). Having found no evidence that the association between offspring survival and impending maternal death was explained by the mother’s chronological age or by differences in offspring exposure to extrinsic mortality risk, we report the results in Figure 2 and Table 1 (which do not account for maternal age or differences in extrinsic mortality risk) as our final results.

### *Second prediction:* F1 offspring that experience maternal loss will produce F2 offspring with compromised survival

To test for intergenerational effects of early maternal loss (Figure 1, gold arrow), we built species-level mixed effects Cox proportional hazards models of offspring survival until the end of a species-specific immature period. The predictor of interest in these models is a binary fixed effect indicator of whether the F1 female had experienced maternal loss when she was young, i.e., whether the mother of F1 (M generation, the grandmother of F2) died before the F1 female (the mother of F2) had reached a species-specific age (Table S1). We included random effects of maternal ID as well as birth year at each study site to control for site-specific cohort effects. We also built a capuchin model that combined data from both populations, along with a fixed effect of study site ID.

Three species (muriquis, blue monkeys, and baboons) showed statistically significant intergenerational effects of early maternal loss on offspring survival, such that F1 females who experienced early maternal loss produced F2 offspring who were more likely to die during their immature period as compared to F2 offspring whose mothers did not experience early maternal loss (muriquis: HR = 2.55, p = 0.03; blue monkeys: HR = 1.48, p = 0.049; baboons: HR = 1.28, p = 0.03). Overall, chimpanzees showed the strongest estimated effect, although this relationship did not meet the threshold for statistical significance (HR = 3.1, p = 0.06), likely due to relatively small sample size (n = 7) of F2 offspring having been born to F1 mothers that experienced early maternal loss. Orphaned F1 females chimpanzees reach sexual maturity at a later age(31), and the small sample of F2 chimpanzees may be the result of this and other orphaning effects on F1 reproduction or fertility. No species showed a strong or statistically significant association in the unexpected direction (Table 2, see Figure S2 for species-level survival curves).

**Table 2.** Model outputs of mixed effects models of offspring survival throughout the pre-dispersal immature period as a function of whether the mother of the offspring experienced early maternal loss. All models included site-specific random effects of birth year and maternal ID. Bold values refer to a statistically significant effect (‘p value’ < 0.05) or an estimate in the expected direction (‘In expected direction?’). Italics indicate an estimate where 0.05 < ‘p value’ < 0.1.

| Species | Coef. Estimate | Std. Error | z value | p value | Hazard Ratio Est. [95% Conf. Int.] | In expected direction? |
| --- | --- | --- | --- | --- | --- | --- |
| Chimpanzees | 1.13 | 0.60 | 1.88 | *0.06* | 3.10 [0.96,10.0] | Yes |
| Muriquis | 0.94 | 0.44 | 2.05 | **0.03** | 2.55 [1.07,6.05] | Yes |
| Blue Monkeys | 0.39 | 0.20 | 1.97 | **0.049** | 1.48 [1.00,2.19] | Yes |
| Gorillas | 0.28 | 0.62 | 0.45 | 0.65 | 1.33 [0.39,4.46] | Yes |
| Baboons | 0.25 | 0.12 | 2.14 | **0.03** | 1.28 [1.01,1.62] | Yes |
| Capuchins (Santa Rosa) | -0.32 | 0.35 | -0.92 | 0.36 | 0.73 [0.37,1.44] | No |
| Capuchins (Lomas Barbudal) | 0.13 | 0.27 | 0.48 | 0.63 | 1.14 [0.67,1.92] | Yes |
| Capuchins (Combined)* | -0.07 | 0.21 | -0.32 | 0.75 | 0.93 [0.62,1.41] | No |
| Sifakas | -0.29 | 0.75 | -1.28 | 0.20 | 0.75 [0.17,3.25] | No |

\*Model also included fixed effect of study ID.

Thus far, our approach has not accounted for the potential role of the F1 female’s death on F2 survival in the species that display an intergenerational effect of early maternal loss. It is possible that the observed intergenerational effect of early maternal death operates exclusively by increasing the likelihood that F1 will die while F2 is young, followed by F2 death thereafter. We tested this possibility in the three species that displayed intergenerational effects of early maternal loss on offspring survival.

First, in blue monkeys and baboons, we added a day-by-day time-varying binary indicator of the F1 female’s presence to our survival model of the F2 offspring’s survival throughout the immature period. This approach required that we exclude those cases in which F2 and F1 death dates were identical (N=58 F2 baboons and N=3 F2 blue monkeys). For blue monkeys, the estimate of the magnitude of the intergenerational effect was nearly identical (0.396 vs 0.393), although the estimate no longer met the threshold for statistical significance (p = 0.051). Still, the fact that the coefficient estimate was entirely unchanged suggests that the intergenerational effect in blue monkeys was not explained by an increase in the likelihood that F1 would die during F2’s early life.

For baboons, the intergenerational effect that is independent of this F1 death parameter did not meet the threshold for statistical significance (p = 0.08) and the coefficient estimate was somewhat reduced in magnitude (0.21 vs 0.25) as compared to the total intergenerational effect presented in Table 2. This lack of a statistically significant, independent intergenerational effect is likely explained by the strong effect of maternal loss (the M generation) on the adult lifespan of F1 females (26) (Figure 1, red arrow), which in turn increases the chance that F2 offspring will directly experience early maternal loss. However, previous results from the same baboon population indicated a strong and significant intergenerational effect of maternal loss on offspring survival that was independent of maternal death in the offspring generation (p = 0.009, coefficient = 0.34)(33). This previous analysis used a more restricted dataset of offspring and identified 4 years of age as the end of the immature period (as compared to 4.5 years in this analysis, which was selected for consistency with criteria for the other species, see supplement for details).

For muriquis, a formal statistical approach was limited by the small sample size, as only six F2 offspring whose F1 mothers experienced early maternal loss died while they were immature. Of those six individuals, five died while their mothers were still alive, and those mothers did not die in the near future, indicating that any intergenerational effect of early maternal loss is independent of any effect on F1 death. In the sixth muriqui case, uncertainty in death dates creates uncertainty as to whether the F2 or F1 individual died first. Regardless of the order of events in this final case, the data from muriquis suggest that the intergenerational effect of maternal loss cannot be explained by an increased likelihood of death in the F1 generation.

Taking these results together we conclude that the intergenerational effects of early maternal loss on offspring survival in these three species are largely independent of the offspring’s direct experience of maternal presence or absence, although the two are perhaps not completely independent in baboons.

## Discussion

We have shown that in several primate species, offspring face reduced fitness prospects if they are born to mothers that will die in the near future (i.e., within a few years) or if they are born to mothers that themselves had experienced early maternal loss. Our first analysis shows that, in five species, offspring whose mothers will die in the near future face reduced fitness prospects *before* maternal death actually occurs, presumably because they experience reductions in maternal allocation (Figure 2, Table 1). This negative association between impending maternal death and offspring survival is also strong and significant when data from all seven species are considered together. Furthermore, in three of the species considered, the effects of early maternal loss carry over from one generation to the next (Figure 3, Table 2), as offspring are less likely to survive the immature period if their mothers had experienced early maternal loss. We interpret this intergenerational effect to be the result of chronic reductions in the physical condition of female offspring that experience early maternal loss.

### Why are offspring more likely to die when their mothers will die in the near future?

The result that offspring exhibited reduced survival in years 0-2 if their mother was going to die in the near future is consistent with our predictions. We propose that offspring survival and future maternal death are both affected by an unmeasured variable: the physical condition of the mother in the years shortly after offspring birth. Our interpretation is that mothers who die are often in poor physical condition for several years before their death, and that this reduction in condition has negative effects on their offspring, presumably through reductions in the quantity or quality of maternal care (hereafter the “maternal condition hypothesis”). This interpretation is also consistent with findings from humans that young children experience reduced survival if their mother is critically ill (38).

However, an alternative hypothesis to explain the observed association between offspring survival and future maternal death is that offspring death during the earliest stage of its life has a causal effect on future maternal death as a result of increased psychological stress experienced by the mother that accompanies the loss of an offspring (hereafter the “maternal grief hypothesis”). Studies of humans has found that parents who experience the death of a child face reduced health and survival for years afterward (39–44), including an increase in the likelihood of deaths associated with grief (e.g., deaths by suicide (45–47)).

### Distinguishing between the maternal condition and maternal grief hypotheses

We therefore seek to distinguish between the maternal condition hypothesis and the maternal grief hypothesis to the extent that we are able. Some of the data regarding the causes of death for infant chimpanzees are generally consistent with the maternal condition hypothesis. Of the 13 infant chimpanzees who died within the first two years of life and whose mothers died shortly thereafter, 3 were cases in which an offspring was born to an SIV-positive mother, who died shortly after her offspring died. In a fourth case, the mother was known to be very ill with a different disease at the time of the offspring’s death. These cases are consistent with similar data from humans regarding survival of infants born to HIV positive mothers(38), which implicate poor maternal health in offspring death.

The impact of grief on wild animal behavior and survival are much more challenging to identify in natural populations as compared to humans, but the last decade has seen efforts to identify these effects (48–50). Within the populations studied here, an example of the death of a white-faced capuchin mother from the Lomas Barbudal population is consistent with our expectations of what grief might look like in a wild primate. This female (Chaos) had lost three infants in less than three years, with infanticide suspected to be the cause in each case. In the weeks following the final infant death, Chaos wandered the forest on her own, rarely appearing in daily censuses of her social group. She eventually died, less than five weeks after the death of the third infant. At the same time, Chaos faced additional social challenges, including the deaths of her mother and sister and her immigration into a new social group. So, while we cannot state with certainty that grief resulting from the violent killings of her last three infants contributed to Chaos’ death, her case is fully consistent with the maternal grief hypothesis.

Chaos’ death represents a single case, but it underscores the importance of distinguishing between the maternal condition and maternal grief hypotheses. While Chaos’ death following the killing of her infants is suggestive of maternal grief, other cases of infanticide followed by maternal death could be consistent with the maternal condition hypothesis, if mothers in poor condition are less able to protect their offspring from infanticidal threats. Thus, observations of infanticide followed by maternal death could be consistent with either hypothesis.

We are optimistic that the maternal condition and maternal grief hypotheses can be distinguished through the collection of longitudinal data on maternal care. Specifically, if we assume that the quantity and/or quality of maternal care that a female provides is condition-dependent, then the collection of data about maternal care (e.g. frequency of carrying/nursing infant, nutritional quality of the milk, maternal intervention/protection during dangerous episodes) provided by mothers to offspring born before and after the birth of another offspring that died as an infant would allow for us to distinguish between these two hypotheses. As an example, imagine three offspring that are born to a mother (Figure 4). The first offspring (Offspring A) survives to adulthood, while the second offspring (Offspring B) does not. The third offspring (Offspring C) is born after the death of Offspring B, and the mother dies at some point shortly after the birth of Offspring C. If the maternal grief hypothesis is correct, we would expect the mother to provide equivalent care to offspring A and B, but then provide worse care to Offspring C, as her condition declines due to grief she experiences following the death of Offspring B. Alternatively, if the maternal condition hypothesis is correct, we would expect the mother to provide worse maternal care to Offspring B as compared to Offspring A as she comes closer to death. If her condition continues to decline, we would then expect her to provide even worse care to Offspring C than she provided to Offspring B (see Figure 4). Collection of maternal care data to test these predictions should be carried out concurrently with collection of high-density measures of maternal biomarkers, which could provide a more direct measure of maternal physical condition.

**Figure 4.**
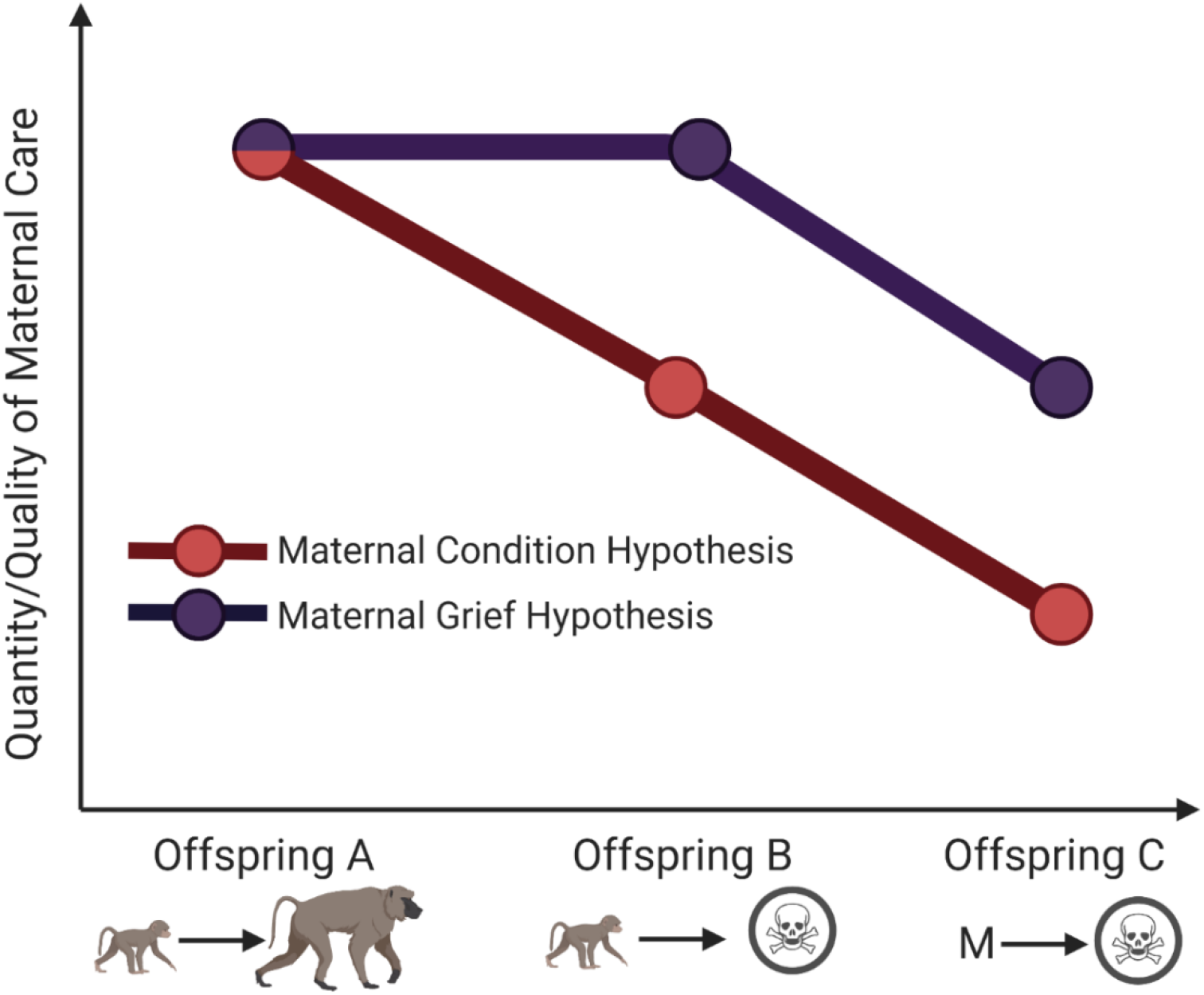
Alternative predictions of the maternal condition and maternal grief hypotheses. In this example, Offspring A survives to adulthood, while Offspring B does not. Offspring C is born after the death of Offspring B, and the mother dies at some point shortly after the birth of Offspring C. Under the maternal grief hypothesis, we predict that maternal care should decline only *after* the death of Offspring B. Under the maternal condition hypothesis, we predict that maternal care should decline *before* the death of Offspring B.

### Variation in the intergenerational effects of early maternal loss

Some, but not all, of the primate species that we considered showed evidence of intergenerational effects of early maternal loss. What might explain this variation? Although a formal phylogenetic analysis will require data from many more species, one possibility is that the intergenerational effects of maternal loss may be reduced in species that display higher levels of allomaternal care. If the mother of the F1 individual dies when F1 is young, but F1 continues to receive care from other relatives (e.g. grandmothers or aunts), this allomaternal care might mitigate the long-term effects of early maternal loss. Simultaneously, allomaternal care provided to the F2 generation could further mitigate any negative effects of reductions in F1 condition. White-faced capuchins displayed one of the lowest coefficient estimates for the intergenerational effect, and are also known to display relatively high levels of allomaternal care as compared to the other species considered here (51–53). Any causal interpretation of this result would be highly speculative at this time, but future work should consider the potential role of allomaternal care in mitigating the costs of maternal loss in wild primates.

### Implications for the evolution of life histories

Our results have important implications for our understanding of the evolution of primate life histories. We have shown that an offspring’s fitness is influenced by the longevity of its mother in ways not previously considered. If we assume that maternal longevity is at least partially under genetic control, then maternal longevity may have an indirect genetic effect on offspring survival and reproductive success (54, 55). Because mothers and offspring are closely related, it follows that mutations that improve maternal longevity would increase females’ fitness not only by extending their reproductive lifespan, but also by decreasing the likelihood that their offspring will experience the detrimental effects of maternal loss during development. We would therefore expect stronger selection on longevity in species in which maternal survival has downstream effects on offspring survival and reproductive success (12, 56, 57). If mothers face an evolutionary tradeoff in which more frequent reproduction leads to reduced lifespan (58), we would expect the associations that we present here to select for a slower, longer life history in female primates as compared to species where links between maternal survival and offspring fitness are weaker or non-existent. Rigorously evaluating such a possibility would require a more formal theoretical approach that is beyond the scope of this paper, but primates’ relatively slow life histories are consistent with this hypothesis (58–60).

We hope that our results encourage other researchers to publish similar analyses that will allow for a broader comparative examination of predictors of whether these effects occur. For example, we predict that all effects of maternal loss will be reduced in species in which allomaternal care is relatively common and important for offspring survival, as other kin and non-kin members of the social group may mitigate the loss of maternal allocation. We also predict that these associations will be strongest in mammal species in which offspring continue to receive support from their mothers beyond the age of weaning, as is the case for most primate species analyzed here. For example, we would expect these associations to be strong in species where mothers and offspring co-reside, such as hyenas and elephants, and less strong in species in which mothers are likely to disperse before the offspring matures, such as gorillas. Further, we expect these associations (especially the blue and purple arrows in Figure 1) to be present across a wide range of the mammalian taxonomy.

More generally, our results reveal unexpected and additional ways in which offspring fitness depends on maternal condition and have important implications for our understanding of life history evolution. Additional studies by behavioral ecologists will be important to confirm if these results are generalizable to other mammal species. Further, our results would benefit from theoretical studies of parent-offspring interactions(61) that can help characterize the microevolutionary dynamics of intergenerational fitness effects.

## Methods

For this study, we analyzed the lives of all offspring included in the Primate Life Histories Database (PLHD; see Results) that met particular inclusion criteria, which we present separately below for the two major analyses that we performed.

### Inclusion Criteria for Analysis #1: Association Between Offspring Death and Impending Maternal Death

To be included in this analysis, three pieces of information needed to be known with certainty: (1) if and when the offspring died prior to age two years, (2) whether the mother of the offspring died within four years of the offspring’s birth, and (3) in cases where the mother died within four years of the offspring’s birth, whether she died before or after her offspring. For mothers and offspring that died on the same day, we make the conservative assumption that the mother died before the offspring. Because we are interested only in offspring survival during the period when the mother is alive, we right-censored the lives of any offspring whose mothers died before them, assigning them a censor date equal to the maternal death date. Offspring were also right-censored at age 2 years, if they survived beyond age 2. The age cutoff of two years is meant to capture a period of life that corresponds to a period of heightened offspring vulnerability for all species considered here. Although the seven species vary in life history patterns, such that some offspring are weaned before two years of age and others are not, offspring of all species are most vulnerable during these first two years of life.

Within the database, many individuals have estimated birth and death dates, with some uncertainty around the dates. If uncertainty in the estimates of demographic events created any uncertainty in the inclusion criteria, those records were excluded from analysis. Table S1 contains the number of offspring included in and excluded from this analysis for each species as well as the number of offspring deaths that contribute to each analysis.

### Inclusion Criteria for Analysis #2: Intergenerational Effects of Early Maternal Loss on Immature Survival

To be included in this analysis, two pieces of information needed to be known with certainty: (1) if and when the offspring (F2 generation) died prior to the end of the species-specific immature period (described in Table S1) and (2) whether the mother (F1 generation) of the F2 offspring had experienced early maternal loss while she was immature (i.e. whether the grandmother, M generation, of the F2 offspring died while the F1 female was immature). As above, if uncertainty in the dates of any demographic events caused ambiguity in any of these areas, records were excluded from analysis. The numbers of offspring included in and excluded from this second analysis also appear in Table S1.

To determine whether the F1 female had experienced early maternal loss, it was first necessary to calculate species-specific estimates of the end of the immature period. For two species (baboons and chimpanzees), we used the median age at which females achieve menarche, which can be directly observed via the appearance of sexual swellings(31, 62). For sifakas, capuchins, and blue monkeys, menarche is not directly observable. We instead estimated the end of the immature period as the median age of first birth, minus one standard deviation(63–65). For gorillas, we estimated the end of the immature period as the median age of first birth, minus two years of “adolescent sterility”(66). Finally, for muriquis, we used the approximate median age of female dispersal the end of the immature period(67). Each of these species-specific estimates of the end of the immature period appear in Table S1.

### Statistical Analyses

For both analyses, we built mixed effects Cox proportional hazards models using the R 3.5.0(68) packages survival(69) and coxme(70).

For all models built for analysis #1, the response variable was time elapsed between offspring birth and offspring death or censoring during the first two years of life. The only fixed effect was a binary indicator of whether the mother died within four years of the offspring’s birth. We included random effects of maternal ID and birth year to control for cohort effects in particular years for each species. For baboons, chimpanzees, and gorillas, we considered the birth year to begin Jan 1 of each calendar year. Birth years for the other species began after the period that included the fewest births(71) (muriquis: March 1; blue monkeys: November 1; sifakas: July 15; capuchins: October 1). We also built combined species models that contained data from all 7 species (or all species except baboons, see results) along with an additional random effect of study site.

For all models built for analysis #2, the response variable was time elapsed between F2 birth and F2 death or censoring during the species-specific immature period (Table S1). The only fixed effect was a binary indicator of whether the F1 female who was the mother of the F2 offspring experienced maternal loss/death when she was immature. As in analysis #1, we also included random effects of maternal ID and birth year at each site in the models.

For those species in which we identified a significant intergenerational effect, we also built post-hoc models that additionally included a time-varying fixed effect of maternal presence in the offspring’s life. This addition allowed us to explore the extent to which the intergenerational effect of early maternal death on F2 survival is independent of an increased likelihood of F_1_ absence during F_2_ early life (i.e. to what extent the gold and red arrows in Figure 1 are independent). All models were assessed for violations of the proportional hazards assumption, and were generally found not to violate the assumption (with a single exception, see Supplement for details).

## Acknowledgments

The Primate Life Histories Working Group was funded by the National Evolutionary Synthesis Center (NESCent), the National Center for Ecological Analysis and Synthesis (NCEAS), the Princeton Centers for Health and Wellbeing and the Demography of Aging, and the Princeton Environmental Institute. The governments of Brazil, Costa Rica, Kenya, Madagascar, Rwanda, and Tanzania provided permission to conduct the field studies. We thank the many students, colleagues, field assistants, and technicians who have contributed to collecting and managing the long-term data stored in the PLHD. We also thank Fernando Colchero for providing helpful comments that improved the manuscript. Specific acknowledgments for each study, including funding agencies, can be found at https://docs.google.com/document/d/1EXHwz473oYgYnuazjixRf40dzltTt0U0maY-vM7vH94/edit?usp=sharing

## Supplementary tables, figures, and materials

**Table S1.**
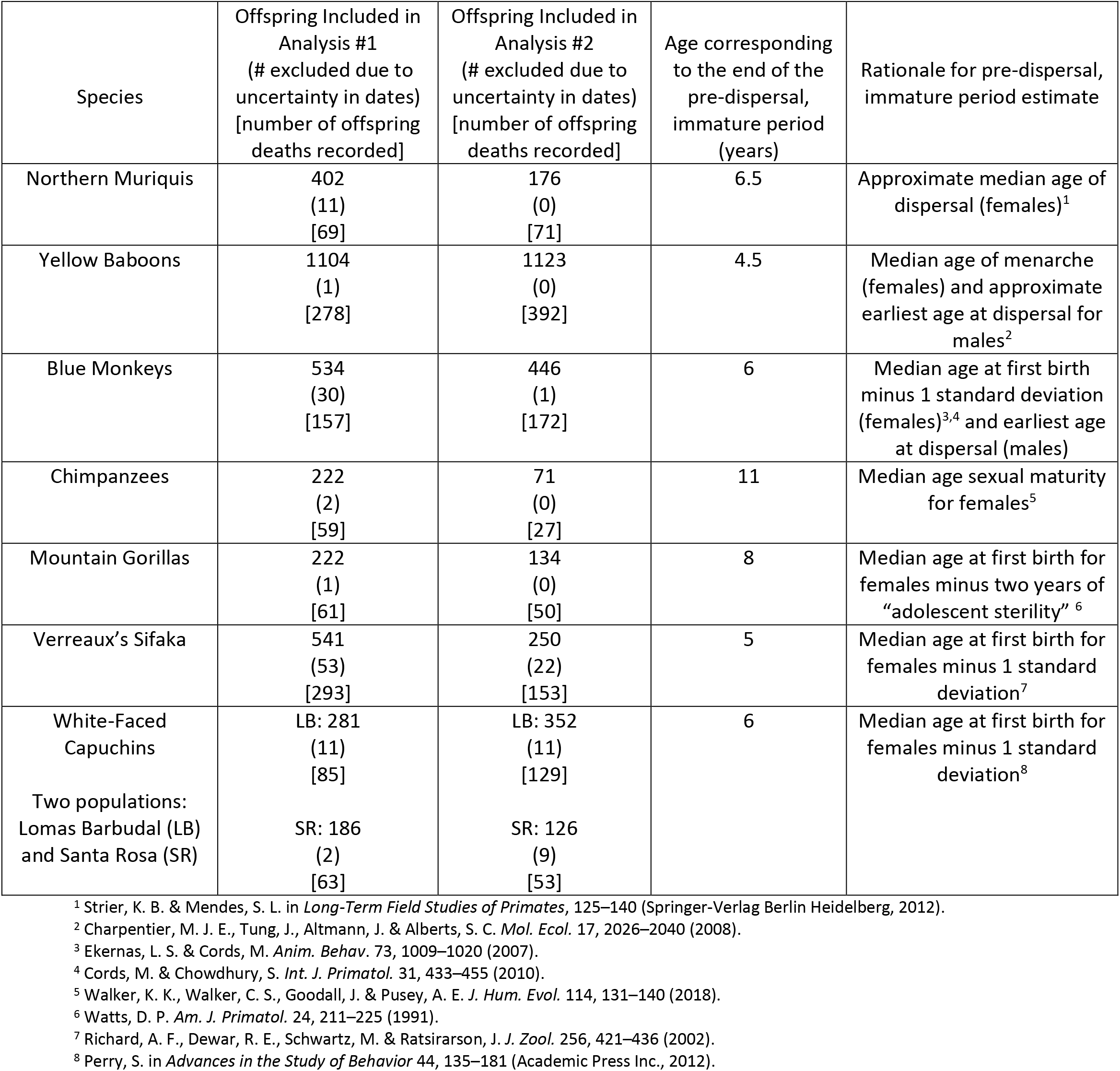
The number of offspring included in each analysis, and estimates of species-specific ages that correspond to the end of the pre-dispersal, immature period (analysis #2).

**Table S2:** Results from mixed effects Cox proportional hazards models of offspring survival to age 2 as predicted by impending maternal death that included an additional term of maternal age, standardized across species. Bold indicates a statistically significant effect (p < 0.05), italics indicate estimates where 0.05< p <0.10.

| Species | Parameter | Estimate | Hazard Ratio<br>[95% Confidence Interval] | p value |
| --- | --- | --- | --- | --- |
| All 7 Species | Impending maternal Death | 0.32 | 0.08 | **< 0.0001** |
|  | Maternal Age (z score) | 0.03 | 0.03 | 0.36 |
| Yellow Baboons | Impending maternal Death | 0.30 | 1.35<br>[1.03,1.78] | **0.03** |
|  | Maternal Age (z score) | -0.015 | 0.98<br>[0.88,1.11] | 0.85 |
| Chimpanzees | Impending maternal Death | 0.80 | 2.22<br>[1.89,4.17] | **0.01** |
|  | Maternal Age (z score) | -0.03 | 0.97<br>[0.75,1.25] | 0.82 |
| Northern Muriquis | Impending maternal Death | 0.72 | 2.05<br>[0.99,4.22] | *0.051* |
|  | Maternal Age (z score) | 0.15 | 1.16<br>[0.91,1.48] | 0.23 |
| Verreaux's Sifaka | Impending maternal Death | 0.26 | 1.29<br>[0.96,1.72] | *8* |
|  | Maternal Age (z score) | 0.04 | 1.04<br>[0.92,1.18] | 0.46 |
| White-Faced Capuchins (Combined) | Impending maternal Death | 0.53 | 1.70<br>[1.17,2.47] | **0.006** |
|  | Maternal Age (z score) | -0.07 | 0.94<br>[0.79,1.11] | 0.44 |

**Table S3.** Results of mixed effects Cox proportional hazards models that predict offspring survival in years 0-2 as a function of impending maternal death. These model results are distinguished from those in Table 1 in the main text by the difference in random effects structure. The models in Table 1 contain study site-specific random effects of birth year, while these models contain social-group specific random effects of birth year. All models in Table 1 and here contain random effects of maternal ID. Bold values refer to a statistically significant effect (‘p value’ < 0.05) or an estimate in the expected direction (‘In expected direction?’ is ‘Yes’). Italics indicate 0.05 < ‘p value’ < 0.10.

| Species | Coef. Estimate | Std. Error | z value | p value | Hazard Ratio Est. [95% Conf. Int.] | In expected direction? |
| --- | --- | --- | --- | --- | --- | --- |
| All Species Combined<sup>^</sup> | 0.34 | 0.08 | 4.11 | **<0.0001** | 1.40<br>[1.20,1.64] | Yes |
| All Species, Except Baboons<sup>^</sup> | 0.39 | 0.10 | 3.77 | **0.0002** | 1.47<br>[1.21,1.80] | Yes |
| Northern Muriquis | 0.80 | 0.36 | 2.21 | **0.03** | 2.22<br>[1.09,4.52] | Yes |
| Chimpanzees | 0.82 | 0.33 | 2.52 | **0.01** | 2.27<br>[1.20,4.28] | Yes |
| White-Faced Capuchins (Santa Rosa) | 0.45 | 0.32 | 1.44 | 0.15 | 1.57<br>[0.85,2.91] | Yes |
| White-Faced Capuchins (Lomas Barbudal) | 0.69 | 0.27 | 2.52 | **0.01** | 2.00<br>[1.17,3.38] | Yes |
| White-Faced Capuchins (Combined)<sup>^</sup> | 0.59 | 0.20 | 2.89 | **0.004** | 1.80<br>[1.21,2.67] | Yes |
| Baboons | 0.28 | 0.14 | 2.04 | **0.04** | 1.32<br>[1.01,1.72] | Yes |
| Sifaka | 0.29 | 0.15 | 1.96 | *0.051* | 1.34<br>[1.00,1.80] | Yes |
| Gorillas | 0.25 | 0.55 | 0.45 | 0.66 | 1.28<br>[0.43,3.79] | Yes |
| Blue Monkeys | 0.03 | 0.24 | 0.12 | 0.90 | 1.03<br>[0.64,1.64] | Yes |

**Figure S1.**
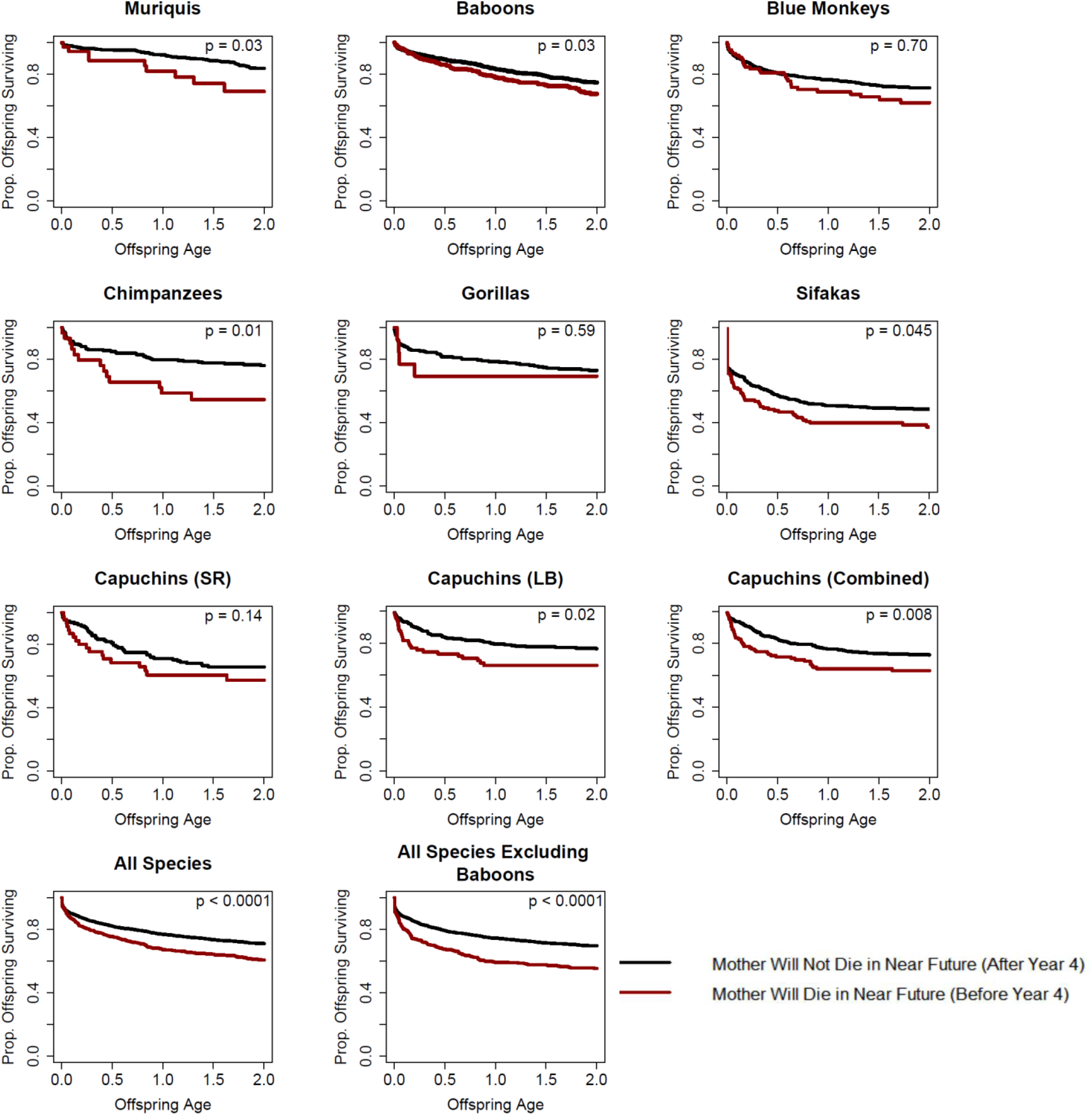
Offspring survival as a function of age and impending maternal death for each of the seven species as well as for all species combined and all species except for baboons. P values refer to the output of a mixed effects cox proportional hazards model of offspring survival through the first two years of life that includes random effects of maternal ID and site-specific birth year. Note that the he combined species models and the combined capuchin models additionally include a random effect of study site ID.

**Figure S2.**
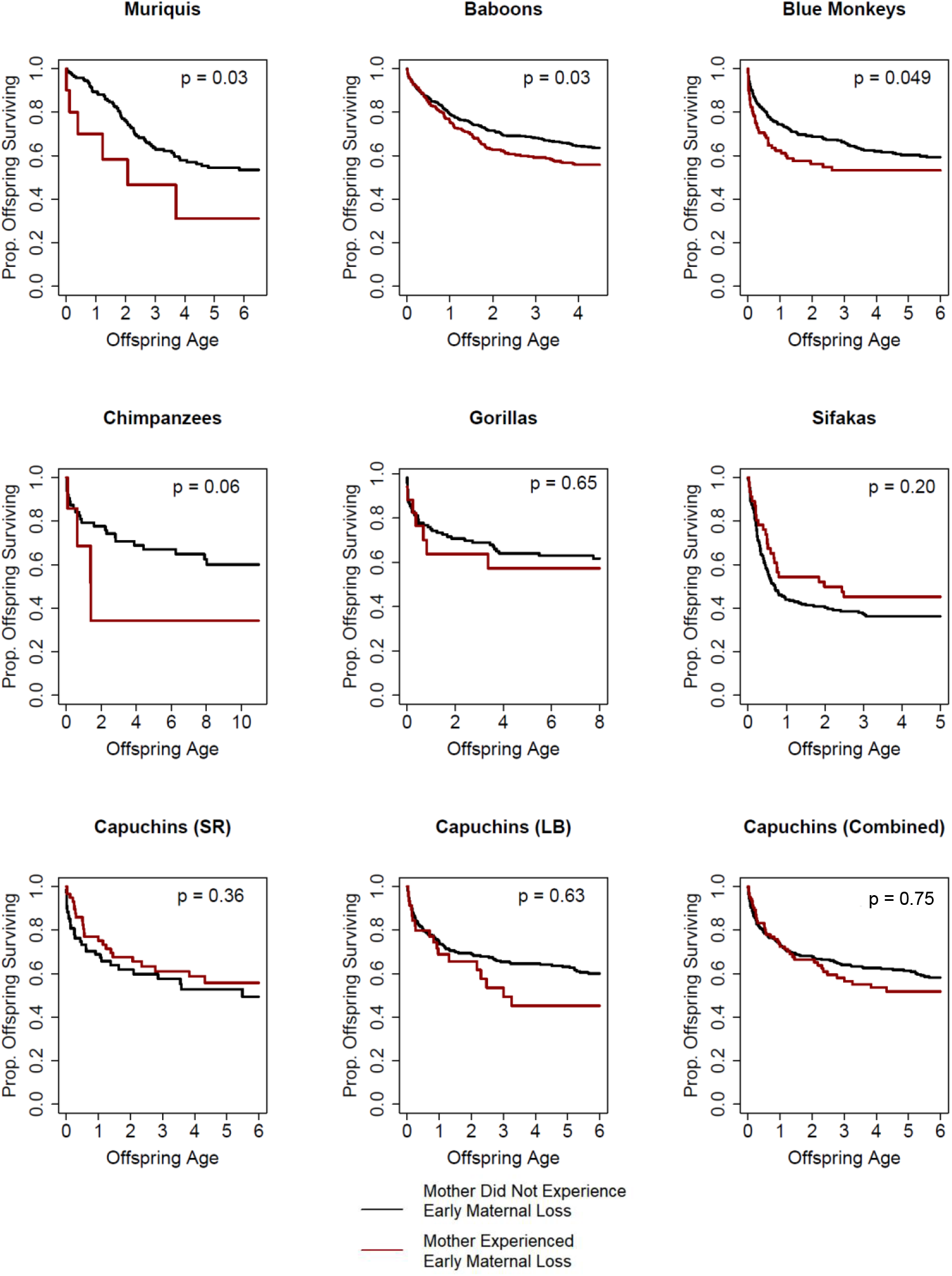
Survival curves for offspring of each species depending on whether the mother of the offspring experienced early maternal loss. P values refer to the output of a mixed effects Cox proportional hazards model of offspring survival throughout the immature period that includes random effects of maternal ID and site-specific birth year. Note that the combined capuchin model also includes a random effect of study site ID.

**Figure S3.**
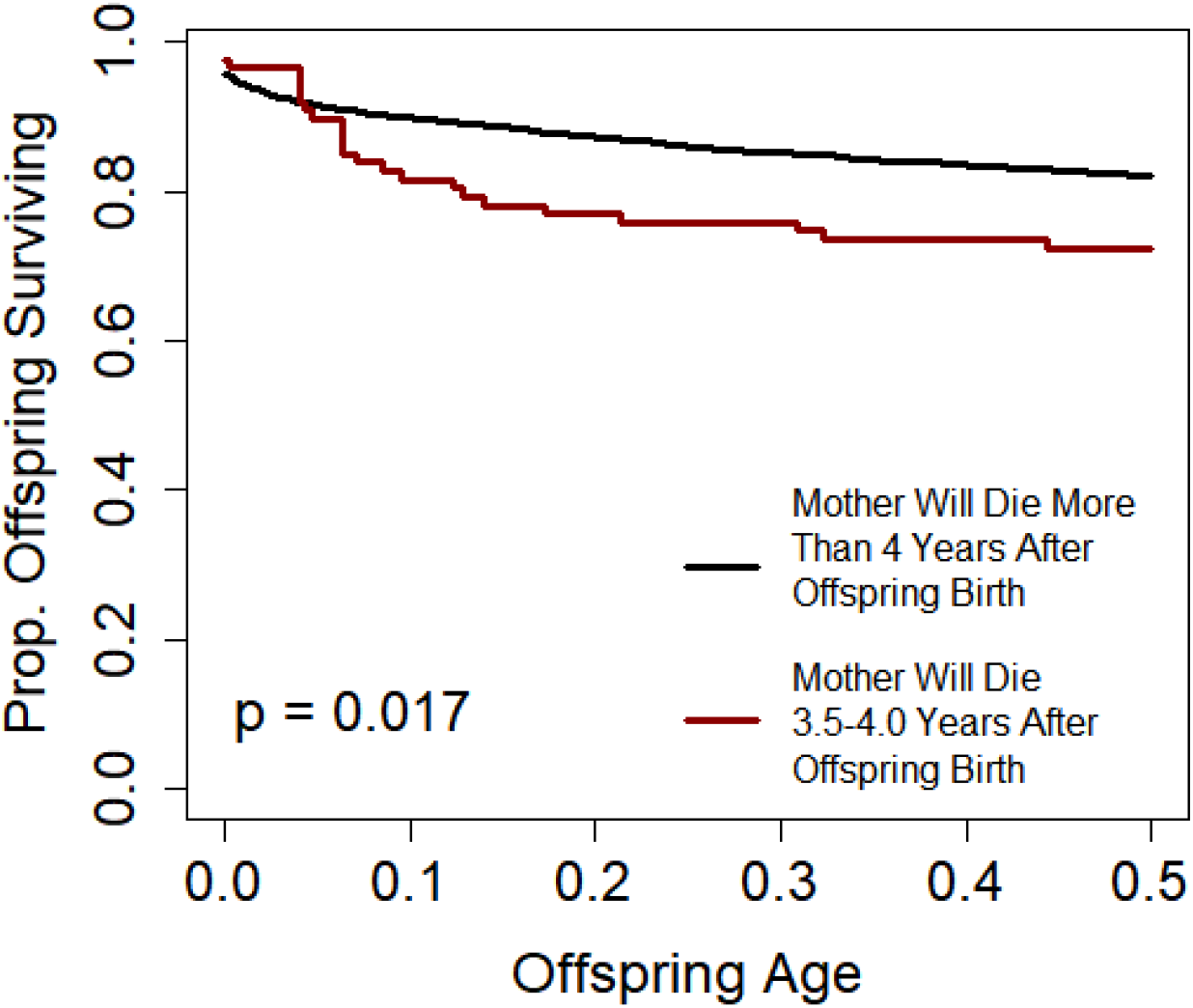
Offspring are more likely to die in the first six months of their life if their mother is going to die in years 3.5-4.0 after the offspring’s birth. This figure contains data from all 7 species, including 84 total offspring whose mothers died 3.5-4.0 years after offspring birth. The displayed p value refers to the output of a mixed effects Cox proportional hazards model of offspring survival during the first six months of life that includes random effects of Maternal ID, study site ID, and site-specific birth year.

### Explanation of differences between baboon dataset in this study and a previous analysis of the intergenerational effect of early maternal loss

A previous study (Zipple *et al* 2019) measured the intergenerational effect of early maternal loss in the Amboseli baboons and identified a strong intergenerational effect that was independent of maternal loss (i.e. death of the F_1_ individual) experienced by the offspring (the F_2_ individual) directly. Our baboon-specific results presented here are on the whole quite similar to those described by Zipple *et al* (2019), but are not identical because we used somewhat different datasets and a different assessment of the end of the baboon immature period in this study than did Zipple *et al* (2019).

Specifically, Zipple *et al* (2019) was a study of the intergenerational effect of multiple sources of early life adversity, not just early maternal loss. For an offspring to be included in that analysis, the authors needed to be able to measure five sources of early adversity, experienced both by offspring directly and by their mothers. For example, it was necessary to know whether the offspring was born to a low-ranking mother and if the offspring’s *mother* had been born to a low-ranking mother. The analytical approach in the present study did not employ such a restriction. Furthermore, Zipple *et al* (2019) restricted their analysis to offspring who had no uncertainty at all about their birth date. We did not impose this stringent requirement in the present study, in part because such a restriction would greatly reduce the sample size in non-baboon species. The end result of these differences in inclusion criteria was that the present study included a larger number of offspring in the intergenerational analysis (n = 1123) than the analysis in Zipple *et al* (2019), which included 687 offspring.

Additionally, Zipple *et al* (2019) analyzed offspring survival in the first 4 years of baboon life, while the present study considers offspring survival to age 4.5. The former study used 4 years as the period of analysis to guarantee that only the immature periods of animals’ lives were being considered (the earliest ages of menarche and natal dispersal is ~4 years in the Amboseli population). In the present study, our estimates of the age corresponding to end of the immature period are meant to represent the median experience of animals in each species, so we used 4.5 years (the median age of menarche in Amboseli) as our estimate of the end of the immature period for baboons. Together, these differences between Zipple *et al* (2019) and the present study likely account for the minor differences in results presented in each study. Both studies are fully consistent with the hypothesis that maternal input is critical a critical determinant of offspring fitness, and that cessation of that input during early life has strong acute and chronic effects.

### Assessing violations of the proportional hazards assumption

We assessed all models contained in Tables 1 and 2 (20 total models) for potential violations of the proportional hazards assumption using the cox.zph function in the *survival* package. Because cox.zph supports coxph objects (rather than coxme objects), we converted our models to coxph objects with frailty terms rather than random effects. Because the coxph function supports only one frailty term per model, we built alternate versions of each model that included each random effect in turn. Across our 20 models, we found evidence that the proportional hazards assumption was significantly violated in only a single case (p < 0.03 for the blue monkey intergenerational analysis; see Table 2).

If we assume that this single case represents a real age-related relationship rather than a type I error, how might it have affected our results? In this case the hazard ratio estimate for offspring whose mothers experienced maternal loss declined significantly as immature offspring aged (see Figure S2, as the separation between red and black lines was greatest during the first two years of life). Therefore, if any bias was introduced into our results by this violation of the proportional hazards assumption, it would have been in the direction of returning a *lower* hazard ratio estimate on average across the immature period as compared to the true effect during the earliest period of offspring life. This violation of assumptions is therefore unlikely to affect our qualitative results, and to the extent that it does, it does so in a *conservative* direction.

